# Cortical Network Disruption is Minimal in Early Stages of Psychosis

**DOI:** 10.1101/2023.12.02.569728

**Authors:** Peter C. Van Dyken, Michael MacKinley, Ali R. Khan, Lena Palaniyappan

## Abstract

1

**Background and Hypothesis:** Chronic schizophrenia is associated with white matter disruption and topological reorganization of cortical connectivity but the trajectory of these changes over the disease course are poorly understood. Current white matter studies in first-episode psychosis (FEP) patients using diffusion magnetic resonance imaging (dMRI) suggest such disruption may be detectable at the onset of psychosis, but specific results vary widely and few reports have contextualized their findings with direct comparison to chronic patients. Here, we test the hypothesis that structural changes are not a significant feature of early psychosis.

**Study Design:** Diffusion and T1-weighted 7T MR scans were obtained from N=113 (61 FEP patients, 37 controls, 15 chronic patients) recruited from an established cohort in London, Ontario. Voxel- and network-based analyses were used to detect changes in diffusion microstructural parameters. Graph theory metrics were used to probe changes in the cortical network hierarchy and to assess the vulnerability of hub regions to disruption. Experiments were replicated with N=167 (111 patients, 56 controls) from the Human Connectome Project - Early Psychosis (HCP-EP) dataset.

**Study Results:** Widespread microstructural changes were found in chronic patients, but changes in FEP patients were minimal. Unlike chronic patients, no appreciable topological changes in the cortical network were observed in FEP patients. These results were replicated in the early psychosis patients of the HCP-EP datasets, which were indistinguishable from controls on nearly all metrics.

**Conclusions:** The white matter structural changes observed in chronic schizophrenia are not a prominent feature in the early stages of this illness.

## 2 Introduction

The neuropathology of schizophrenia comprises a generalized dysconnectivity between brain regions.^1,2^ The current consensus suggests a loss or “subtle randomization” of functional relationships across the brain,^3–7^ predominantly affecting highly connected neural hubs. This causes alterations of the cortical hierarchy,^5,8^ hampering the integrated processing of information and causing disorganization of thoughts, speech, and behaviour.^9,10^

White matter pathology has long been explored as a causative or mediating factor of schizophrenia.^10^ Two decades of diffusion magnetic resonance imaging (dMRI) research, including a mega-analysis of 1963 individuals with schizophrenia,^11^ have established the disruption of white matter (WM) integrity as a robust feature of chronic schizophrenia.^2,12–15^ Although the histological implications of these findings are not entirely clear, they may involve some combination of the degradation of myelin sheaths on long axonal projections,^16^ decreased axonal density^17^ or increased fibre disorganization.^18^

If these changes arise early in the course of schizophrenia, they may reflect causative, pathological processes, and their precise quantification may aid early detection. Such an idea is easily motivated by our current developmental understanding of the disease.^19,20^ A substantial genetic component^21^ and associations with childhood^22^ and perinatal trauma^23^ suggest schizophrenia arises from pathological processes beginning very early in life. Indeed, the first psychotic episode is generally preceded by a subthreshold phase of disorganized thinking and perceptual, motor and cognitive disruptions that can last for several years.^24^ We thus might expect structural changes to have accumulated by the time of the first psychotic episode. If so, such changes may be detectable, early, neural markers of the prodromal period.

Previous dMRI studies in first-episode psychosis (FEP) have thus far converged on a report of reduced fractional anisotropy (FA) in FEP,^25^ but the location and scale of these findings vary across studies.^26–37^ Other reports have studied changes in the number of streamlines connecting cortical regions, a metric that gives insight into the anatomical makeup of the structural connectome. Results from such analyses have been relatively modest compared to FA. Reductions and elevations of streamline counts are observed in scattered connections^38–40^ and the connectivity of cortical hubs is slightly reduced.^41,42^

Common to all the above studies is the relative paucity of findings compared to those observed in chronic patients. In studies analyzing both groups together, disruption is consistently greater in older, more chronic patients compared to those with FEP.^39,43,44^ These prior studies, however, are limited either by limited methodologies^43^ or small sample sizes,^39,44^ especially of FEP patients (*n* <= 20).

The lack of a consistent, anatomically localized disruption in FEP may mask a more reproducible topological effect. Individual deficits may be anatomically scattered, reflecting high inter-individual variability difficult to observe at the group-level, yet still produce a converging effect on the overall topology.^8,45^ Disruption in chronic patients is already known to be topologically biased, with highly connected hub nodes bearing the greatest burden.^12,46^ Previous work has mostly focused on anatomically localized disruption, and no prior studies of FEP patients have studied both anatomical and topological disruption in concert.

Finally, the use of single-site dMRI datasets hinders attempts at replication, as findings may reflect the unique acquisition parameters of the dataset.^47^ The availability of high quality, open-access datasets allows us to replicate observations to ensure robustness of both positive and negative findings. The recently released Human Connectome Project - Early Psychosis (HCP-EP) dataset represents the first such openly available dataset specifically for early psychosis (<3 years since diagnosis). Its diffusion data has not yet been analyzed in any major connectivity study of early psychosis.

In this study, we analyzed geometric and topological disruption in the brains of FEP and early psychosis (EP) patients using the Tracking Outcomes in Psychosis (TOPSY) dataset, a 7T dMRI dataset of untreated FEP patients, age matched controls, and chronic patients, and the HCP-EP dataset. We hypothesize that, compared to chronic patients, FEP and EP patients will have relatively intact network topology and a reduced extent of network disruption.

## 3 Methods

### 3.1 Data

#### 3.1.1 TOPSY Dataset

61 FEP patients, 15 chronic patients, and 37 healthy controls (HCs) were recruited from an established cohort enrolled in the Prevention and Early Intervention Program for Psychoses (PEPP) in London, Ontario. All participants provided written, informed consent prior to participation as per approval provided by the Western University Health Sciences Research Ethics Board, London, Ontario. Inclusion criteria for study participation were as follows: for FEP patients: individuals experiencing first-episode psychosis, with no more than 14 days of cumulative lifetime antipsychotic exposure, no major head injuries (leading to a significant period of unconsciousness or seizures), no known neurological disorders, and no concurrent substance use disorder. Participants were not explicitly instructed to abstain from substances, and patients on non-antipsychotic prescription medication were not excluded.

For FEP, the mean lifetime total defined daily dose days (DDD × days on medication) for antipsychotic use was 1.57 days with 27 patients (47.4%) being completely antipsychotic naive at the time of scanning. Of those who had started antipsychotic treatment, (N = 30; 52.6%), the median total defined daily dose days was 2.81 days (range of 0.4–14 DDD days). Patients with established (chronic) schizophrenia consisted of clinically stable patients on long-acting injectable medications with 3 or more years since illness onset, no recorded hospitalization in the past year, and receiving community-based care from physicians affiliated to a first-episode clinic (PEPP, London Ontario). While many studies have focused on chronic schizophrenia patients in their 40s or 50s, our approach enabled us to reduce (even if we cannot fully avoid) the age gap with FEP and HC groups. Patient consensus diagnosis was established using the best estimate procedure described by Leckman *et al.*^48^ following 6 months of treatment. Diagnoses were made based on the Structured Clinical Interview for DSM-5.

HCs were recruited through posters and word-of-mouth advertising. Healthy control subjects had no personal history of mental illness, no current use of medications, and no first-degree relatives with a history of psychotic disorders. Healthy controls were group matched to the FEP cohort for age and parental socioeconomic status (the National Statistics Socioeconomic Classification: five-class version). Similar to their FEP counterparts, those with a history of substance use disorders in the past 12 months, significant head injury, or neurological disorders were excluded.

Data was acquired with a head-only, neuro-optimized 7T MRI (Siemens MAGNETOM Plus, Erlangen, Germany) using dMRI and T1-weighted (T1w) imaging protocols. T1w data was collected using an MP2RAGE sequence^49^ at 0.75 mm isotropic resolution, echo time = 2.83 ms, repetition time = 6 s, field of view = 240×240 mm, number of slices = 208. The T1w image was reconstructed using the robust algorithm introduce by O’Brien *et al*.^50^ Diffusion data was acquired with an echo-planar imaging (EPI) sequence at 2mm isotropic resolution, echo time = 50.2 ms, repetition time = 5.1 s, field of view = 208 mm, number of slices = 72, MB acceleration factor = 2, flip angle = 90. 64 directions were acquired in both the AP and PA directions at b=1000, along with 2 b=0 images. Gradient nonlinearity correction was applied to all acquisitions using in-house software.

#### 3.1.2 HCP-EP Dataset

Human Connectome Project - Early Psychosis (HCP-EP) data was accessed according to the Data Use Certification issued by the NIMH Data Archive. Data was acquired on a 3-T MRI (Siemens MAGNETOM Prisma). T1w data was collected using an MPRAGE sequence at 0.8mm isotropic resolution, echo time = 2.22ms, repetition time = 2.4 s, field of view = 256 mm, number of slices = 208. dMRI was acquired with an EPI sequence at 1.5mm isotropic resolution, echo time = 89.2 ms, repetition time = 3.23 s, field of view = 210 mm, number of slices = 92, MB acceleration factor = 2, flip angle = 78. 92 directions were acquired in both the AP and PA directions at b=1500 and b=3000, along with 7 b=0 images.

### 3.2 Preprocessing

#### 3.2.1 Anatomical data

For TOPSY data, segmentation of the anatomical images and construction of the cortical surface mesh was performed using *FastSurfer*,^51–53^ a recently developed implementation of the cortical parcellation and mesh creation algorithms pioneered by *FreeSurfer*, chosen for its improved processing efficiency and more accurate parcellations (as determined with visual quality control). Remaining processing of anatomical images was done using *ciftify*,^54^ an implementation of the HCP-EP minimal preprocessing workflow.^55^ Of note, images were registered to the *MNI152NLin6Asym*^56^ template space, and meshes were registered to the *fsLR-32k* template space.^57^

For the HCP-EP dataset, we used the minimally preprocessed anatomical images included in the dataset.^55^

#### 3.2.2 Diffusion Data

Diffusion data was preprocessed using *snakedwi*,^58^ a preprocessing pipeline based on *snakebids*^59^ and *snakemake*.^60^ Briefly, Gibbs ringing artefacts were removed with *mrdegibbs* from *MRtrix3*;^61,62^ eddy currents and motion were corrected using *eddy* from *FSL*;^63^ susceptibility-induced distortions were corrected using *topup* in *FSL*, using the AP-PA pairs of images.^64,65^ The T1w image was skull stripped with *SynthStrip*;^66^ bias field correction was applied with *N4ITK* from *ANTS*.^67^ A T1 proxy image was created from the diffusion image using *SynthSR*^68^ and used to accurately register the diffusion image space to the T1w space using *greedy*.^69^ Diffusion tensor imaging (DTI) metrics were calculated using *dtifit* from *FSL* with linear regression.^70^

Tractography was performed using the *MRtrix3*^62^ software suite. Constrained Spherical Deconvolution (CSD) was performed using the dhollander algorithm to estimate the response functions for WM, grey matter (GM) and cerebrospinal fluid (CSF).^71,72^ Single shell 3 tissue CSD (SS3T-CSD), as implemented in *MRtrix3Tissue* (https://3tissue.github.io/), was performed to obtain WM-like fibre orientation distibutions (FODs) as well as GM-like and CSF-like compartments in all voxels.^73^ mtnormalise was used to correct for residual intensity inhomogeneities.^74,75^ Tractography was performed using the iFOD2 algorithm^76^ and anatomically constrained tractography (ACT),^77^ with 50,000,000 streamlines, an FOD amplitude cut-off of 0.06, a minimum streamline length of 4mm, and a maximum streamline length of 250mm. The anatomical segmentation used for ACT was obtained using *SynthSeg*.^78^ Spherical-deconvolution informed filtering of tractograms 2 (SIFT2) was used to correct the streamline counts based on the underlying FOD magnitude.^79^

The HCP-EP dataset was preprocessed in the same manner, except that multi-tissue CSD^80^ was used instead of SS3T-CSD, and 10,000,000 streamlines were generated in tractography. All b-values were used for the calculation of DTI metrics.

### 3.3 Analysis

Voxelwise differences in FA, mean diffusivity (MD), radial diffusivity (RD), and axial diffusivity (AD) were calculated using tract-based spatial statistics (TBSS).^81^ DTI metrics were used because more complicated models could not be fit to our TOPSY dataset, which has a maximum b-value of 1000. FA maps were first registered to a common template space roughly corresponding to the average of all FA images using a pipeline based on *greedy*.^69^ The other DTI maps were then transformed into this space. *FSL*^65^ was then used to skeletonize the maps, and *FSL* `randomise`^82^ with TFCE-correction and 10,000 permutations was used to compare groups, thresholding at a corrected p-value of 0.05. Sex and age were regressed as nuisance variables. For HCP-EP data, acquisition site was additionally regressed.

Subject anatomical scans were parcellated using the Brainnetome atlas.^83^ These parcellations were used to derive weighted connectivity matrices based on the tractography data. Average FA, MD, RD, AD, and the log-transformed SIFT2-weighted streamline count were used as weights. Edges in the resulting graphs with significantly differing weights across groups were identified using network based statistic (NBS)^84^ with extent based cluster sizes, a T threshold of 3.0, 10,000 iterations, and corrected p-value threshold of 0.05. Sex and age (and site for HCP-EP) were regressed as nuisance variables.

Node hubness was calculated using a composite score^85^ based on four graph theory metrics: degree, betweenness, clustering coefficient, and path length, all of which have been previously used to capture hubness.^86^ These parameters were defined according the weighted definitions given by van den Heuvel et al.^86^ These parameters were calculated for each node, and the nodes were rank-ordered and assigned a score based on their rank. The score was defined as the node’s position in the rank-ordered list divided by the total number of nodes. In other words, each node had a score between 0 and 1, with 0 given to the node with the lowest value, and 1 given to the node with the highest. Such a score was calculated independently for each of the four metrics, and the four scores for each node were averaged to get the overall hubness score. This entire process was repeated independently for each subject, yielding subject-specific hubness scores for all nodes. For group analyses, these hubness scores were averaged across the subjects within each group.

Differences in node hubness were analyzed by comparing the relative rank order of nodes between and within groups. First, Kendall’s tau^87^ was used to compare hubness rankings for all subject pairs. This metric varies between −1 and 1, where 1 means the two lists have the same order, 0 means the two lists have uncorrelated orders, and −1 means the two are reversed relative to each other. We then calculated the average within-group and between-group Kendall’s tau, i.e. the average metric for all subjects in one group compared to all subjects within the same group, and the average metric for all subjects in one group compared to all subjects in another group. Significance was assessed by randomly permuting the groups 10,000 times.

Connection disruption and node hubness were compared based on an analysis described in,^12^ with some modification. To measure the disruption associated with nodes of a given hubness, nodes were selected from the hubness band surrounding threshold *k* ± *r* where the kernel radius *r* was set to 0.05. The proportion of disrupted edges connected to these nodes was compared to 10,000 randomly selected groups of nodes of equal size. This analysis was repeated at values 0.1 < *k* < 0.9. This analysis was repeated using the ranked degree of each node, where each node was assigned a value between 0 and 1 based on its position in a degree-ranked list of nodes. Finally, both of these analyses were repeated using a threshold approach. Here, *k* was treated as an upper or lower threshold. For each value of *k*, the subgraph of nodes above or below *k* was considered, and only edges within that subgraph were evaluated. Empirical disruption proportions were compared to 1000 equivalently sized, randomly selected subgraphs.

### 3.4 Code Availability

The analyses discussed above and resulting figures were made possible by openly available python packages,^88–98^ particularly *graph-tool*^99^, *pybids*,^100,101^ *nibabel*,^102^ and *nilearn*.^103^ All code used is freely available at https://github.com/pvandyken/paper-CorticalDisruptionFEPMinimal. Links to pipelines used for data preprocessing are listed at that repository.

## 4 Results

### 4.1 Demographics

In the TOPSY dataset, no significant difference was found between HCs and FEP patients for sex, age, handedness, or SES. A similar lack of significant differences were found between HCs and chronic patients, except that chronic patients were significantly older than the HC cohort (*t* = 4.72, *p* < 0.0001). Both FEP and chronic patients had significantly lower education levels, higher cannabis use, and higher smoking prevalence than controls (Table 1).

**Table 1:** TOPSY Demographics: Group distribution columns show *mean* (*sd*) unless otherwise indicated. Comparison columns show *test*(*df*) = *statistic*. P values less than 0.05 are shown in bold. M = male, F = female, SES = Socioeconomic status, CAST = Cannabis Abuse Screening Test, AUDIT-C = Alcohol Use Disorders Identification Test, DUP = Duration of Untreated Psychosis, DUI = Duration of Untreated Illness, SOFAS = Social Occupational Functioning Assessment Scale, PANSS-8 = Positive and Negative Syndrome Scale 8

|  | HC<br>(n=37) | FEP<br>(n=61) | chronic<br>(n=15) | HC vs FEP | HC vs chronic | FEP vs<br>chronic |
| --- | --- | --- | --- | --- | --- | --- |
| Sex (M/F) | 24/13 | 50/11 | 12/3 | $\chi^2(1) = 2.78$,<br>$p = 0.096$ | $\chi^2(1) = 0.547$,<br>$p = 0.46$ | $\chi^2(1) = 0$, $p = 1$ |
| Age | 21.70<br>(3.49) | 22.82<br>(4.65) | 28.93<br>(7.92) | $t(96) = -1.26$,<br>$p = 0.21$ | $t(49) = -4.55$,<br>$p < \mathbf{0.001}$ | $t(73) = -3.83$,<br>$p < \mathbf{0.001}$ |
| Handedness<br>(R/L/A) | 34/0/3 | 53/1/7 | 13/1/1 | $\chi^2(2) = 0.928$, $p = 0.63$ | $\chi^2(2) = 2.53$,<br>$p = 0.28$ | $\chi^2(2) = 1.42$,<br>$p = 0.49$ |
| Education | 14.14<br>(2.18) | 12.83<br>(1.86) | 12.71<br>(2.33) | $t(94) = 3.12$,<br>$p = \mathbf{0.0024}$ | $t(48) = 2.03$,<br>$p = \mathbf{0.047}$ | $t(72) = 0.205$,<br>$p = 0.84$ |
| SES | 3.08<br>(1.34) | 3.53<br>(1.32) | 3.40<br>(1.24) | $t(85) = -1.55$,<br>$p = 0.13$ | $t(49) = -0.785$,<br>$p = 0.44$ | $t(64) = 0.339$,<br>$p = 0.74$ |
| CAST | 6.68<br>(2.77) | 12.14<br>(5.76) | 11.54<br>(7.78) | $t(86) = -5.33$,<br>$p < \mathbf{0.001}$ | $t(48) = -3.3$,<br>$p = \mathbf{0.0018}$ | $t(62) = 0.311$,<br>$p = 0.76$ |
| AUDIT-C | 3.03<br>(2.19) | 2.28<br>(2.97) | 3.38<br>(2.40) | $t(81) = 1.27$,<br>$p = 0.21$ | $t(48) = -0.494$, $p = 0.62$ | $t(57) = -1.23$,<br>$p = 0.23$ |
| Smoker (yes/no) | 1/36 | 17/44 | 7/8 | $\chi^2(1) = 8.12$,<br>$p = \mathbf{0.0044}$ | $\chi^2(1) = 12.6$,<br>$p < \mathbf{0.001}$ | $\chi^2(1) = 1.2$,<br>$p = 0.27$ |
| Cannabis (yes/no) | 10/27 | 36/19 | 5/8 | $\chi^2(1) = 11.6$,<br>$p < \mathbf{0.001}$ | $\chi^2(1) = 0.178$,<br>$p = 0.67$ | $\chi^2(1) = 2.17$,<br>$p = 0.14$ |
| DUP (weeks) -<br>median (IQR) | N/A | 18.50<br>(60.50) | N/A |  |  |  |
| DUI (weeks) -<br>median (IQR) | N/A | 139.00<br>(216.00) | N/A |  |  |  |
| Antipsychotic<br>Duration (days) | N/A | 1.56<br>(3.05) | N/A |  |  |  |
| Antipsychotic<br>Defined Daily<br>Dose Equivalents | N/A | 1.08<br>(3.07) | N/A |  |  |  |
| SOFAS | 82.03<br>(4.67) | 41.62<br>(12.82) | 56.00<br>(11.41) | $t(91) = 17.2$,<br>$p < \mathbf{0.001}$ | $t(45) = 11.2$,<br>$p < \mathbf{0.001}$ | $t(74) = -3.97$,<br>$p < \mathbf{0.001}$ |
| PANSS-8 Total | 8.00<br>(0.00) | 24.96<br>(6.94) | 14.50<br>(7.08) | $t(92) = -14.8$,<br>**$p < 0.001$** | $t(49) = -5.68$,<br>**$p < 0.001$** | $t(69) = 5.03$,<br>**$p < 0.001$** |
| PANSS-8<br>Positive | 3.00<br>(0.00) | 11.95<br>(2.82) | 6.71<br>(3.81) | $t(92) = -19.3$,<br>**$p < 0.001$** | $t(49) = -6.03$,<br>**$p < 0.001$** | $t(69) = 5.79$,<br>**$p < 0.001$** |
| PANSS-8<br>Negative | 3.00<br>(0.00) | 7.71<br>(4.32) | 4.29<br>(1.90) | $t(93) = -6.62$,<br>**$p < 0.001$** | $t(49) = -4.19$,<br>**$p < 0.001$** | $t(70) = 2.89$,<br>**$p = 0.0052$** |
| PANSS-8 General | 2.00<br>(0.00) | 5.34<br>(2.18) | 3.57<br>(2.56) | $t(93) = -9.31$,<br>**$p < 0.001$** | $t(49) = -3.79$,<br>**$p < 0.001$** | $t(70) = 2.64$,<br>**$p = 0.01$** |

In the HCP-EP dataset, no significant differences between HCs and patients were found for sex, SES. Patients were significantly younger (*t* = 3.51, *P* < 0.001), more right-handed (*χ*^2^ = 4.36, *P* < 0.05), less educated (*t* = 3.23, *P* < 0.05), and more likely to smoke (*χ*^2^ = 3.88, *P* < 0.05) than the HCs. Cannabis use was not reported amongst HCs, but 29% of patients had at least some exposure (Table 2).

**Table 2:** HCP-EP Demographics: Group distribution columns show *mean* (*sd*). Comparison columns show *test*(*df*) = *statistic*. P values less than 0.05 are shown in bold. SES = Socioeconomic status, PANSS = Positive and Negative Syndrome Scale

|  | HC (n=56) | Patient (n=111) | HC vs Patient |
| --- | --- | --- | --- |
| Sex (M/F) | 37/19 | 68/43 | $\chi^2(1) = 0.192, p = 0.66$ |
| Education | 15.88 (1.92) | 14.21 (2.20) | $t(73) = 3.23, p = 0.0019$ |
| Age | 24.90 (4.08) | 22.73 (3.59) | $t(165) = 3.51, p < 0.001$ |
| Handedness (R/L) | 43/10 | 99/7 | $\chi^2(1) = 4.36, p = 0.037$ |
| SES | 2.07 (1.07) | 2.25 (1.19) | $t(163) = -0.955, p = 0.34$ |
| Smoke (Yes/No) | 3/51 | 20/89 | $\chi^2(1) = 3.88, p = 0.049$ |
| Pack years | 0.02 (0.09) | 0.71 (2.50) | $t(131) = -1.82, p = 0.071$ |
| Cannabis | N/A | 1.56 (0.89) |  |
| Antipsychotic Duration<br>(months) | N/A | 14.32 (15.65) |  |
| Antipsychotic Defined Daily<br>Dose Equivalents | N/A | 163.51 (231.00) |  |
| PANSS Total | N/A | 49.60 (10.74) |  |
| PANSS Positive | N/A | 11.05 (3.87) |  |
| PANSS Negative | N/A | 14.10 (5.32) |  |
| PANSS Global | N/A | 24.50 (5.26) |  |

### 4.2 Microstructure

We first looked for evidence of global disruption of WM in FEP and chronic patients. No difference was found between the global average FA between FEP patients and HCs, but a significant reduction of FA in chronic patients compared to HCs was noted (*p*_*adj*_ = 0.010). We did not find a difference of FA between the early psychosis patients in the HCP-EP dataset and their associated controls (Figure 1 A).

**Figure 1:**
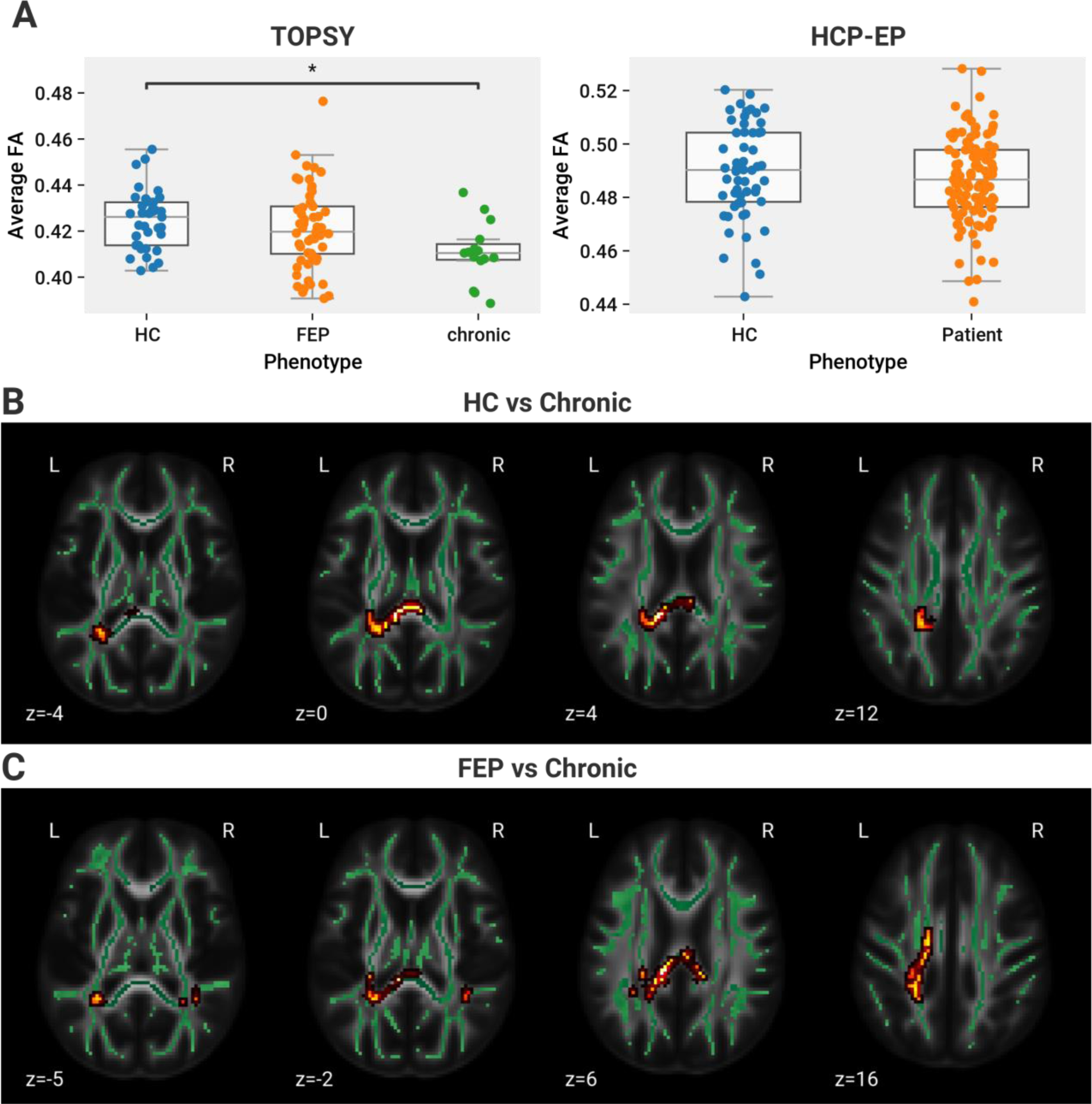
Global FA values are reduced in chronic but not FEP and early psychosis. A: Global average of FA across brainnetome connectome connections. Left side shows data from TOPSY dataset (one-way ANOVA, *F*(2,110) = 4.44, *p* = 0.014); post hoc analysis found a significant reduction of FA in chronic patients compared to HCs (Tukey HSD, FWER=0.05, *p*_*adj*_ = 0.010). A trending difference was observed between FEP and chronic patients (*p*_*adj*_ = 0.0504). Right side shows HCP-EP, no significant difference was found. B,C: Skeletonized voxels with significantly lower FA in chronic patients compared to (B) HCs and (C) FEP patients, as show by FSL’s TBSS. Results as shown are inflated for visualization; the actual significant voxels are restricted to the skeleton, portrayed as a green overlay. Results are thresholded for a FWER of 0.05. Localization and size of the significant voxels is listed in Table S1.

Compared to HCs, chronic patients also had a significant increase in global MD (*p* < 0.001), RD (*p* < 0.001), and AD (*p* = 0.0025). Compared to FEP patients, chronic patients had significantly increased global MD (*p* = 0.0053) and RD (*p* = 0.0034) (Figure S1). No significant differences in these parameters were observed between EP patients and HCs in the HCP-EP dataset.

To narrow down differences to a more regional level, we used TBSS to perform spatial comparisons across HCs, FEP patients, and chronic patients (adjusted for age and sex). No significant differences were found between HCs and FEP. When comparing chronic patients to HCs, voxels with significant FA reduction were found in the left hemisphere occupying the posterior junction of the corpus callosum and corona radiata (Figure 1 B). Further significant voxels were found in chronic patients in comparison to FEP patients, located bilaterally in the posterior corpus callosum and superior longitudinal fasciculus, and in the left hemisphere corona radiata. (Figure 1 C). No significant differences were found between HCs and patients in the HCP-EP dataset.

TBSS performed on other parameters also revealed extensive changes in both chronic and FEP patients. Both patient groups had significantly higher MD than HCs in corpus callosum; the bilateral corona radiata, thalamic radiation, and superior longitudinal fasciculus; and in the superficial white matter of the parietal and occipital lobes (Figure 2 A,C). The MD changes in FEP patients were generally more constrained to the superficial white matter and peripheral white matter tracts, as opposed to the deeper, centralized changes in the chronic patients covering more of the corpus callosum and internal capsule. Chronic patients had higher MD than FEP patients only in the right hemisphere, in the splenium of the corpus callosum and the posterior corona radiata (Figure 2 E). Chronic patients had higher RD than both HCs (Figure 2 B) and FEP patients (Figure 2 F) throughout the corpus callosum, bilaterally in the internal capsule, corona radiata, thalamic radiations, superior longitudinal fasciculus, and tapetum, as well as bilateral superficial white matter in the parietal, temporal, and occipital lobes. Finally, FEP patients had higher AD than HCs in the left hemisphere genu of the corpus callosum, anterior limb of the internal capsule, anterior corona radiata, external capsule, and superficial white matter of the frontal lobe (Figure 2 D). The full list of impacted regions and extent volume is tabulated in Table S1. No significant results were found when testing any of the above models in reverse (e.g. testing for regions in HCs with higher MD than FEP or chronic patients).

**Figure 2.**
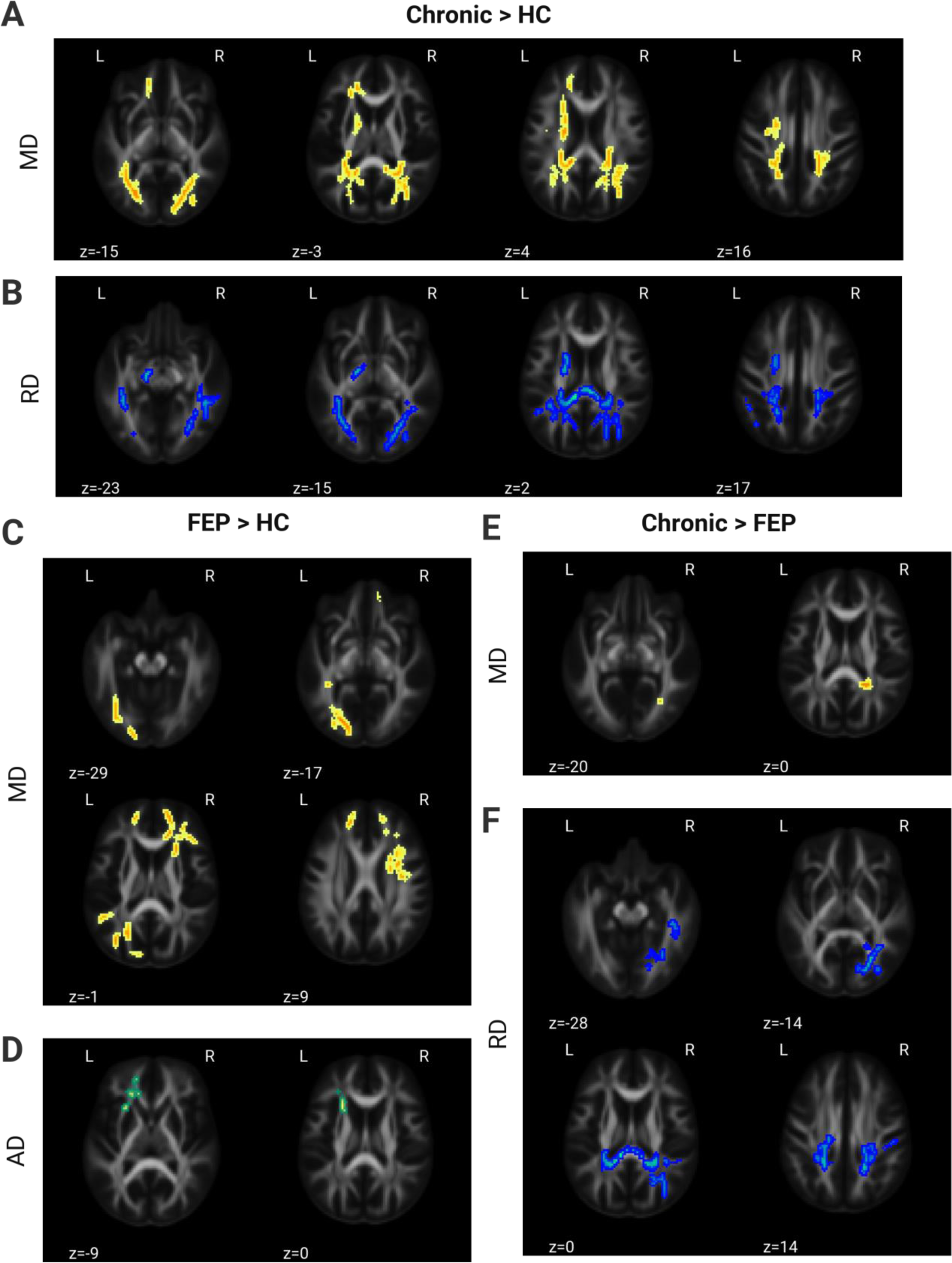
MD and RD increased in chronic patients relative to healthy controls (HCs) and first-episode psychosis (FEP) patients; MD and AD increased in FEP patients relative to controls. All figures show skeletonized voxels with a significantly higher parameter value in the indicated patient group. Results as shown are inflated for visualization, coloured according to the parameter they map (yellow=MD, blue=RD, green=AD), and thresholded for a family-wise error rate of 0.05. The size and localization of significant voxels is listed in Table S1. A and B show results in chronic patients versus healthy controls: A=MD, B=RD. C and D show FEP patients versus healthy controls: C=MD, D=AD. E and F show chronic versus FEP patients: E=MD, F=RD. Abbreviations: MD=Mean Diffusivity, AD=Axial Diffusivity, RD=Radial Diffusivity.

No significant differences of MD, RD, and AD were found between EP patients and HCs in the HCP-EP dataset using TBSS.

### 4.3 Connectivity

To explore the effects of schizophrenia on connectivity, we used the Brainnetome cortical parcellation^83^ and NBS^84^ to search for connections with significantly reduced FA and streamline count, comparing across HCs, FEP patients, and chronic patients. No specific connections with significantly lower FA were found in FEP patients when compared to HCs, but many were found both in chronic patients when compared to HCs or FEP patients (Figure 3 A). The disrupted edges were predominantly interhemispheric, primarily originating from the temporal cortex, insula, and frontal cortex (Figure 3 B). In the HCP-EP dataset, a small subnetwork of disrupted connections restricted to the occipital cortex was found in FEP patients compared to HCs. No differences were found between any group when looking at streamline count-weighted connectomes (Figure 3 A).

**Figure 3:**
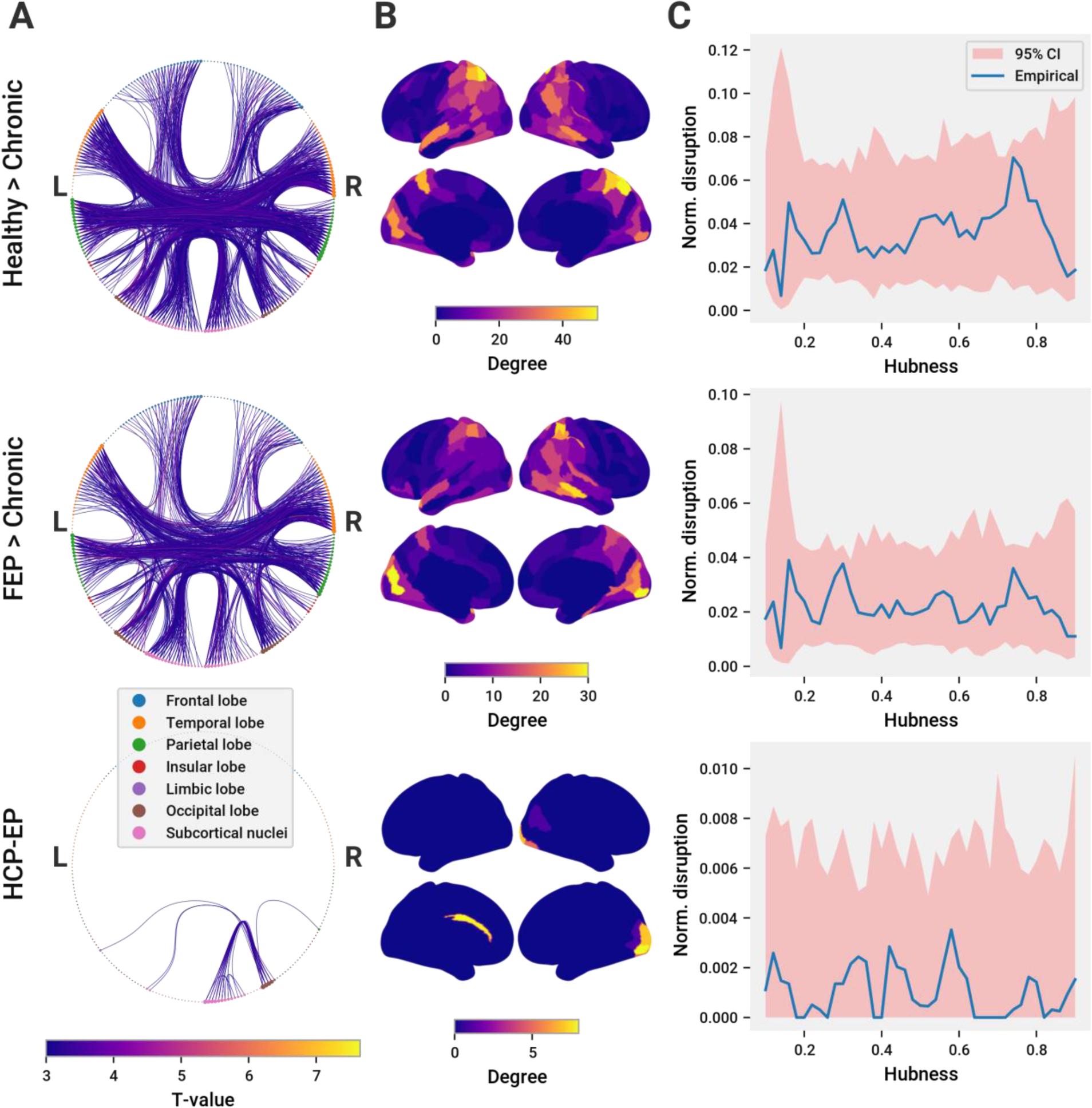
Chronic patients have more disrupted edges than FEP and early psychosis. In all panels, the first row corresponds to results comparining HCs to chronic patients, the second comparing FEPs to chronic patients, and the third comparing HCs in the HCP-EP dataset to the early psychosis patients in that datset. A: Edges with significantly reduced FA as discovered with NBS. Nodes corresponding to the Brainnetome parcellation are represented around the edge of each graph diagram, are coloured according to lobe they belong to, and sized according to the number of disrupted edges they connect with. Left hemisphere nodes are on the left side of the figure, right hemisphere nodes are on the right. Visualized edges are those with significantly lower FA in the control group (FEP in the FEP v chronic comparison). Edges are coloured according the the T-value determined by NBS. B: Spatial distribution of nodes with disrupted edges. Regions are coloured according to the number of disrupted edges connected to the region. C: Hubness of disrupted edges. X-axis corresponds to a band of hubness values corresponding to the hubness *x* ± 0.05. Nodes within that hubness band are selected and the proportion of disrupted edges connected to those nodes is calculated. This proportion is compared to 1000 randomly selected groups of nodes of equal size. The 95% confidence interval of this random distribution is represented by the red shaded region.

Additionally, numerous edges with increased MD, AD, and RD were found in chronic patients compared to both HCs and FEP patients (Figure S2 A,B). A smaller number of edges with increased MD and AD were additionally found in FEP patients compared to HCs (Figure S2, C). Disrupted edges primarily originated from temporal, parietal, occipital, and subcortical regions, and were predominantly interhemispheric. Disrupted edges in FEP patients were predominantly in the frontal lobe and intrahemispheric.

### 4.4 Topology

Changes in the cortical hierarchy have previously been associated with schizophrenia. In particular, studies have found disrupted edges to be predominantly between topologically central nodes. To investigate this, we performed a variation of an analysis previously reported by Klauser et al.,^12^ which measured the proclivity of connections above a given threshold degree to be disrupted, as determined by NBS. Here, we defined a composite metric of hubness comprising four complementary graph theory measures: degree, average shortest path length, betweenness centrality, and the clustering coefficient. All metrics were calculated using the structural connectome weighted with the SIFT2-corrected streamline count. Across all subjects, hubs were situated in locations previously implicated by the literature, including the prefrontal cortex, anterior and posterior cingulate cortex, precuneus, and parietal cortices (Figure 4 D). We measured the probability that edges of a given hubness would be disrupted compared to the general probability of disruption, using a sliding window approach. In all comparisons, there was no particular hubness regime with a disproportionate extent of disruption (Figure 3 C). This finding remained true when using a threshold version of the approach more comparable to the original paper: instead of nodes within a fixed, sliding kernel, all nodes above a sliding threshold were analyzed (Figure S3 A). Finally, no significant results were observed when using an upper-bound threshold instead of a lower-bound (Figure S3 B), and when using the rank-ordered degree of nodes instead of hubness (Figure S3 C,D,E).

**Figure 4.**
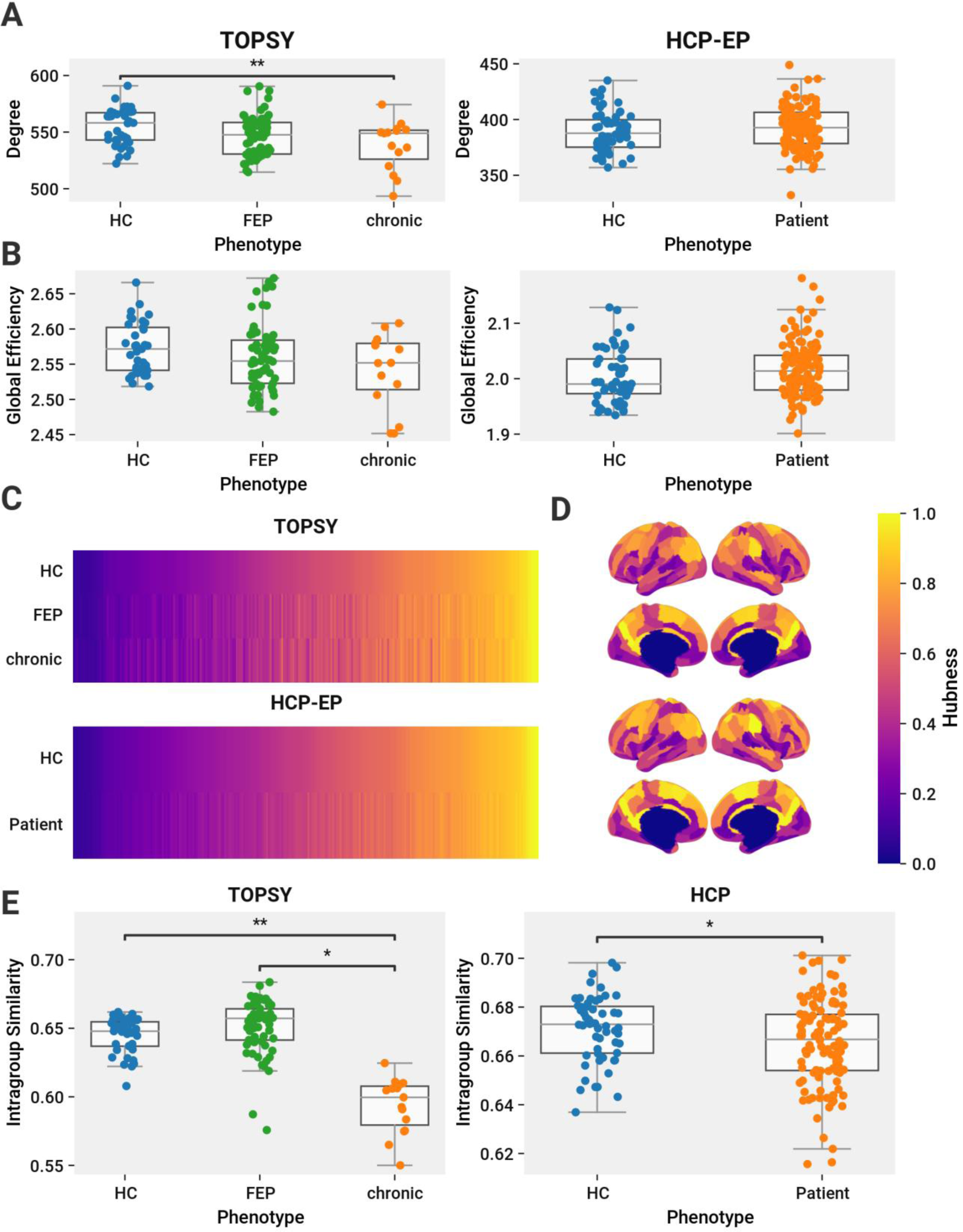
Evidence of topological disruption in chronic but not in FEP or EP patients. A,B: Comparison of graph features calculated on the Brainnetome-parcellated connectome weighted with the logarithm of the SIFT2-weighted streamline count. TOPSY dataset is shown on the left column, HCP-EP on the right. A: Average degree across all nodes. A significant difference was found across all groups in the TOPSY dataset (one-way ANOVA, *F*(2,110) = 4.95, *p* = 0.0088), with post hoc analysis finding a significant reduction in degree in chronic patients compared to HCs (Tukey HSD, FWER=0.05, *p* = 0.0093). No significant difference was found in the HCP-EP dataset. B: Average global efficiency across all nodes. No differences were found between any groups in either the TOPSY or HCP-EP datasets. C: Average rank-order of hub nodes across groups. The top graph corresponds to the TOPSY dataset, the bottom to the HCP-EP. X-axis corresponds to the Brainnetome connectome nodes, rank-ordered for each graph in order of increasing hubness for the datasets’s HCs. Disease conditions are ordered in the same order as HCs for qualitative comparison. D: Spatial distribution of nodes. Each region is coloured according to its hubness as shown in C. E: Rank-order within-group similarity calculated using Kendall’s tau. The left chart displays comparisons within the TOPSY dataset, the rightmost compares HCs and patients in the HCP-EP dataset. Chronic patients have significantly less internal similarity than both HCs (permutations=10,000,) and FEP patients (permutations=10,000,). The right chart show the same analysis for the HCP-EP dataset. EP patients have less internal similarity than HCs (permutations=10000, *p* = 0.034).

To investigate changes in the topology and hierarchy of cortical connectivity, we compared the average degree and global efficiency using the SIFT2-weighted structural connectomes. A significant reduction in average degree was observed between HC and chronic patients (*p* = 0.0093), but not between any of the other groups (Figure 4 A). No changes in global efficiency were found in any comparison (Figure 4 B). Changes in cortical hierarchy across groups were measured by comparing the rank-ordered list of nodes sorted by increasing within-group average hubness (Figure 4 C,D). Similarity was measured using the Kendall Tau Rank correlation coefficient. The hubness rankings of chronic patients has significantly more within-group variation than both HCs (*p* = 0.019) and FEP patients (*p* = 0.035) (Figure 4 E). Furthermore, chronic patients significantly varied from both HCs (*p* = 0.0022) and FEP (*p* = 0.50) patients compared to a permutation-based null distribution (Figure S4 B). No other groups showed differences in hub ranking.

Importantly, the hierarchical changes observed in chronic patients were notably subtle. Although chronic patients show a statistically significant hub reorganization from HCs and FEP patients based on hubness rankings, the overall rankings remain broadly similar across all patients (Figure S4 A). Furthermore, hierarchical disruption in chronic patients cannot be characterized by any stereotyped rearrangements; instead, the individual rankings of patients are increasingly idiosyncratic, as shown by the decreased within-group similarity compared to HCs and FEP patients (Figure 4 E).

### 4.5 Age-matched Healthy Controls and Chronic Patients

To see if the differences between HCs and chronic patients were driven by their difference in age, we repeated significant analyses using an older subset of HCs (n=8) age-matched to our chronic patients. Demographic comparisons between these two groups are shown in Table S2. Differences in global FA (*t*(17) = 2.29, *p* = 0.035) persisted, but no significant differences were found in sift2-based average connection degree (Figure S5 A). Many connections had significantly lower FA as determined by NBS, although not as many as the main comparison (family-wise error rate=0.05) (Figure S5 B,C). No significant clusters were found in the TBSS subgroup analysis.

### 4.6 Duration of Untreated Psychosis

To see if global FA was related to the duration of untreated psychosis (DUP) in FEP, we modified our TBSS and NBS analyses to test for a correlation between FA and DUP in our FEP patients. *N*=47 patients had a valid DUP recorded. The mean DUP was 60.17 weeks, but the median was 21, reflecting a highly left-skewed distribution. We thus took the log of DUP before applying the regression with FA. No clusters were significantly associated with DUP in our TBSS analysis, and no significantly disrupted subnetwork was found with NBS. This analysis was not repeated in the HCP-EP data, as patients in that dataset have a variable duration of treated psychosis, obfuscating any potential effect of DUP.

## 5 Discussion

Focusing on the nature of WM changes in FEP, EP, and chronic schizophrenia, we report 3 key observations: (1) In both untreated FEP and treated early stages of psychosis, structural changes of the WM are minimal, observable only at the level of a few individual tracts (Figure 3), and not at a predictable spatial location (Figure 1); (2) consequently, both global metrics of WM integrity and systems-level topological properties (degree, efficiency, similarity of hub distribution) are unaltered in these early stage samples, when compared to healthy individuals; (3) in contrast, there is a widespread deterioration of tract-level, regional, and systems-level integrity in chronic schizophrenia when compared to healthy subjects and the disease’s early stages. Taken together, these results indicate that WM changes may not be necessary causal factors for the onset of psychosis but may play a role in progression of the illness to chronic stages.

Given the system-wide patterns of disruption observed in chronic patients^12^ and the long prodromal period preceding the first psychotic episode,^24^ one might expect a widespread, accumulated disruption of connectivity in FEP patients. Observation of such disruption would strengthen the case for the primacy of structural dysconnectivity in the clinical expression of psychosis. However, our results demonstrate that connectivity deviations at the resolution measurable by MRI are minimal by the time of the first psychotic episode, eliminating extensive WM structural decline as a necessary precondition for the expression of psychotic symptoms. The pronounced WM disruption seen in established cases of schizophrenia likely reflect secondary mechanisms distinct from the causal pathway of symptom expression.

Nevertheless, there were some connections and regions in FEP patients with significantly higher MD and AD than HC. Notably, a large number of voxels in the right frontal lobe and left posterior lobe had significantly higher MD. Strikingly, although a similar number of voxels had higher MD in both FEP and chronic patients compared to HCs, only a fraction of the connections in FEP patients had higher MD (Figure S2 C.1) compared to the broad effect on connectivity seen in chronic patients (Figure S2 A.1). This illustrates the importance of location in driving network disruption. The MD changes in chronic patients were in central WM regions, around the internal capsule and junctions of the corpus callosum and corona radiata. Changes in these crossroad regions intersect with a vast number of association and commissural tracts, amplifying their effect on the cortical network. Illustrating this effect, only a small anatomical region has higher MD in chronic patients than FEP patients (Figure 2 E), but its central location by the corpus callosum impacts a large number of connections (Figure S2 B.1). On the other hand, the extensive but peripheral MD increases in FEP patients relative to HCs (Figure 2 C) translate to comparatively few disrupted connections (Figure S2 C.1).

FEP patients furthermore maintain the structural hierarchy of the WM network. Across all participants in both datasets, the hub nodes are generally focused in the prefrontal cortex, cingulate gyrus, and precuneus, with high quantitative similarity between individuals (Figure 4 E). While chronic patients all have the same general distribution, they have significantly lower rank similarity compared to other chronic patients, FEP patients, and HCs (Figure S4 B). We note this observation was made in spite of the lack of any particular connections with a significantly different streamline count, reflecting its topological, rather than anatomical, basis.

Thus, both anatomically and topologically, we observe disparities between FEP and chronic patients. Three potential non-causal mechanisms may contribute to this.

First, structural decline may be a secondary phenotype of upstream disease processes. While we lack longitudinal data to conclusively demonstrate this hypothesis, we cite three lines of evidence supporting this notion. First, functional connectivity is known to be disrupted in FEP,^104–106^ and long term interregional signalling patterns modify the strength of the connecting tracts,^107,108^ and can affect tract myelination.^109^ Thus, aberrant functional signalling, over time, may lead to reorganization of the anatomical network. Second, schizophrenia has been associated with neuroinflammation and myelin degradation. For instance, a meta-analysis by Najjar et al found strong evidence for neuroinflammatory pathology in the WM of chronic schizophrenia patients.^110^ Third, several studies of FEP have found aggravated findings resulting from a longer DUP. Kraguljac *et al.* found patients with longer DUP had lower global FA levels.^25^ A more focused study of the tapetum found a similar result.^111^ Note that Filippi *et al.* have reported the opposite trend.^31^ In our own results, we did not observe a relationship between DUP and FA, however, our participants were significantly skewed toward low DUP, limiting our ability to observe an effect. More study of the progression of untreated psychosis is needed to understand the impact of early intervention and the disease halting potential of medication.

A second source of change may be the psychoactive drugs used therapeutically by schizophrenia patients. Since most participants with chronic schizophrenia are undergoing treatment and treatment cannot be ethically withheld, disambiguating the effects of the disease and treatment is generally an ill-posed problem. A few studies, however, have achieved natural cross-sectional experiments. In a rare sample of 17 unmedicated patients with chronic schizophrenia and an age and illness duration matched group of treated patients, Luo *et al.* compared WM integrity^112^ between groups. Unmedicated patients had slightly greater deviation from controls than treated patients, indicating that antipsychotics may not be contributing to the reduced WM integrity *per se*. Another report found no difference in FA reduction between medicated and unmedicated patients.^12^ Thus, evidence to date is more suggestive of antipsychotics either ameliorating or limiting the WM deficits associated with psychosis, rather than contributing to them.

Finally, network disruption may not result from schizophrenia or its treatment at all. Instead, a subset of early-stage patients with lower WM integrity may preferentially progress towards chronic stages of illness. If so, this would introduce a selection bias when recruiting a sample of established cases of schizophrenia. The enrolled patients are likely to be the ones with severe-enough illness that prompted continued engagement with the health care system. Our recruitment approach for the TOPSY dataset selected from a consecutive samples of all referrals to the only first episode program within our catchment, thus including patients irrespective of the later severity and retention probability. Thus, our patient cohort may display a broader array of neurological phenotypes, which, on average, becomes indistinguishable from healthy controls. This would create an opportunity to find subgroups of patients with greater deviation, which might in turn be predictive of long-term outcomes. Such exploration will be a focus of future work.

The healthy controls recruited for our analysis were age-matched to our FEP patients, making age a pertinent difference between both of these groups and our chronic patients. Age has been previously associated with FA decline, however, the onset of this decline is typically observed between 40 and 50 years of age.^43,113–115^ Our chronic patients have a mean age of 30, too early for age related changes to have effect. Accordingly, our post-hoc analysis between chronic patients and an age-matched subgroup of controls still found significant FA reductions in the patients, and NBS found a large number of disrupted edges, although TBSS failed to produce significant findings, possibly due to the smaller sample sizes involved.^28^

Our results stand in contrast to some reports. In chronic schizophrenia, most disrupted structural connections have been observed between the highly connected hub nodes found in the rich club of the brain.^12,86,116^ We did not observe this pattern of disruption between any of our groups. This may be due to our relatively small sample size of chronic patients. A study by Cui et al.^41^ reports a reduction in connectivity amongst the rich-club nodes in FEP, and another found generalized disruption of network efficiency and reported fewer hubs in FEP,^42^ but we were not able to replicate this in either early psychosis dataset. To be sure, our FEP and EP datasets are sufficiently powered and larger than many prior studies that report WM deficits in patient groups. While this may be due to a greater patient heterogeneity discussed above or a study effect stifling “negative” findings such as we report here, we cannot exclude the potential effects of DUP,^25,111^ age, scan parameters, and processing choices in these differences. We note, however, that unlike data used in previous studies, the HCP-EP dataset is freely available for research allowing for future replicability.

Our results do not support the longstanding hypothesis that extensive pre-onset disruption of WM tracts contributes to the development of psychosis. Although some localized diffusivity changes can be observed in FEP patients, these effects are restricted to the periphery and fail to impact global connectivity in the manner observed in chronic patients. When considered with the accumulating evidence discounting prominent grey matter changes preceding first episode psychosis,^117^ these findings suggest reduced WM integrity in schizophrenia may reflect an accumulated burden wrought by severe mental illness over a sufficiently long period of time, rather than an upstream cause of psychotic phenotypes.

## 6 Acknowledgements

Special thanks to Ravi Menon for his leadership in establishing the initial TOPSY cohort.

Research using Human Connectome Project for Early Psychosis (HCP-EP) data reported in this publication was supported by the National Institute of Mental Health of the National Institutes of Health under Award Number U01MH109977. The HCP-EP 1.1 Release data used in this report came from DOI: <u>10.15154/1522899</u>.

