## Supplemental Data for "Cortical Network Disruption is Minimal in Early Stages of Psychosis"

### 1 SUPPLEMENTARY FIGURES

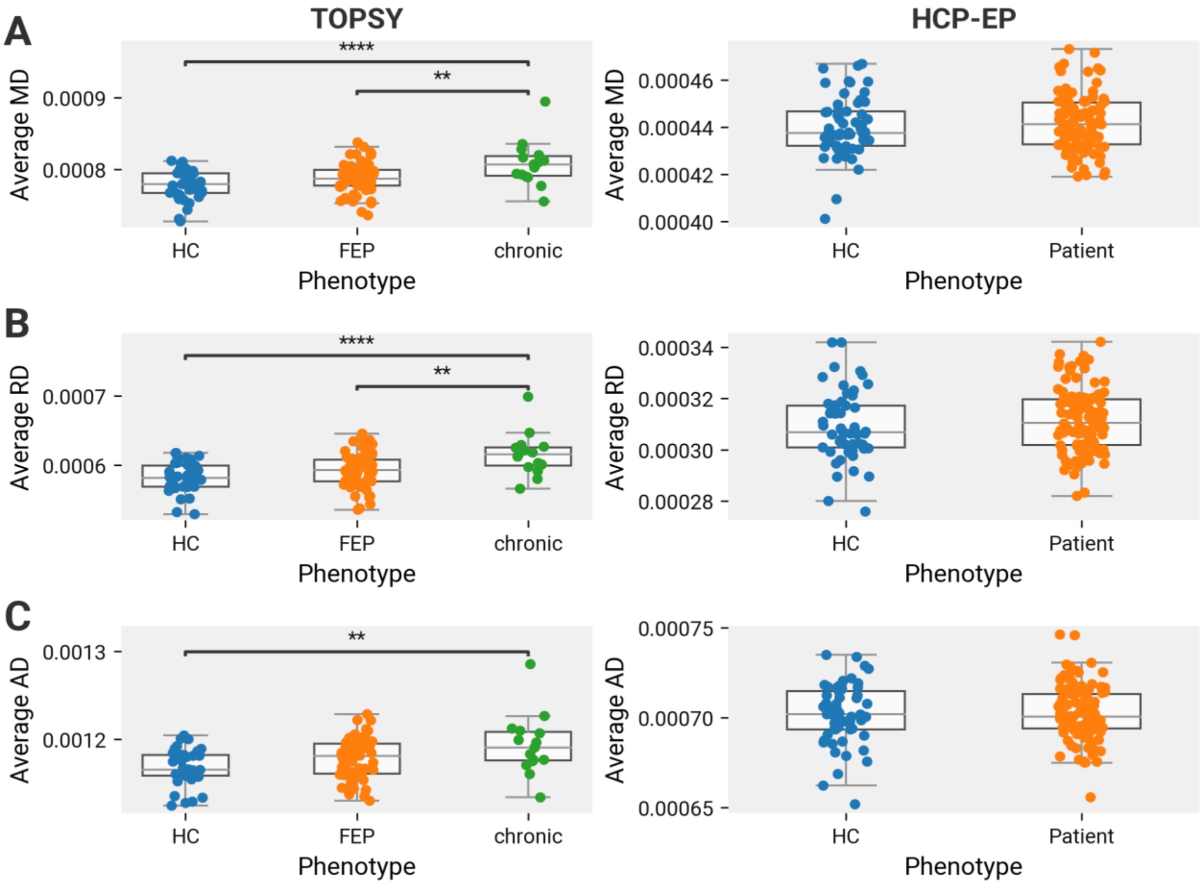


*Figure S1: Chronic, but not first-episode psychosis (FEP) and early psychosis (EP) patients, are different from healthy controls (HCs). Left column shows data from TOPSY dataset, right shows data from HCP-EP. All comparisons within TOPSY are tested using one-way ANOVA with follow-up post hoc analysis using Tukey's HSD with the family-wise error rate controlled to 0.05. All comparisons within HCP-EP are tested using two-sample t-test. No differences were found in HCP-EP in any of the metrics analyzed. A. Average MD. In TOPSY, significant MD differences found between groups ($F(2{,}110) = 9.76, p < 0.001$), with significantly higher MD in chronic patients compared to both HCs ($p < 0.001$) and FEP patients ($p = 0.0053$). B. Average RD. In TOPSY, significant RD differences found between groups ($F(2{,}110) = 9.89, p < 0.001$), with significantly higher RD in chronic patients compared to both HCs ($p < 0.001$) and FEP patients ($p = 0.0034$). C. Average AD. In TOPSY, significant AD differences found between groups ($F(2{,}110) = 6.02, p = 0.0033$), with significantly higher AD in chronic patients compared to HCs ($p = 0.0025$) but not FEP patients. Abbreviations: MD=Mean Diffusivity, RD=Radial Diffusivity, AD=Axial Diffusivity, TOPSY=Treatment Outcomes in Psychosis, HCP-EP=Human Connectome Project - Early Psychosis.*

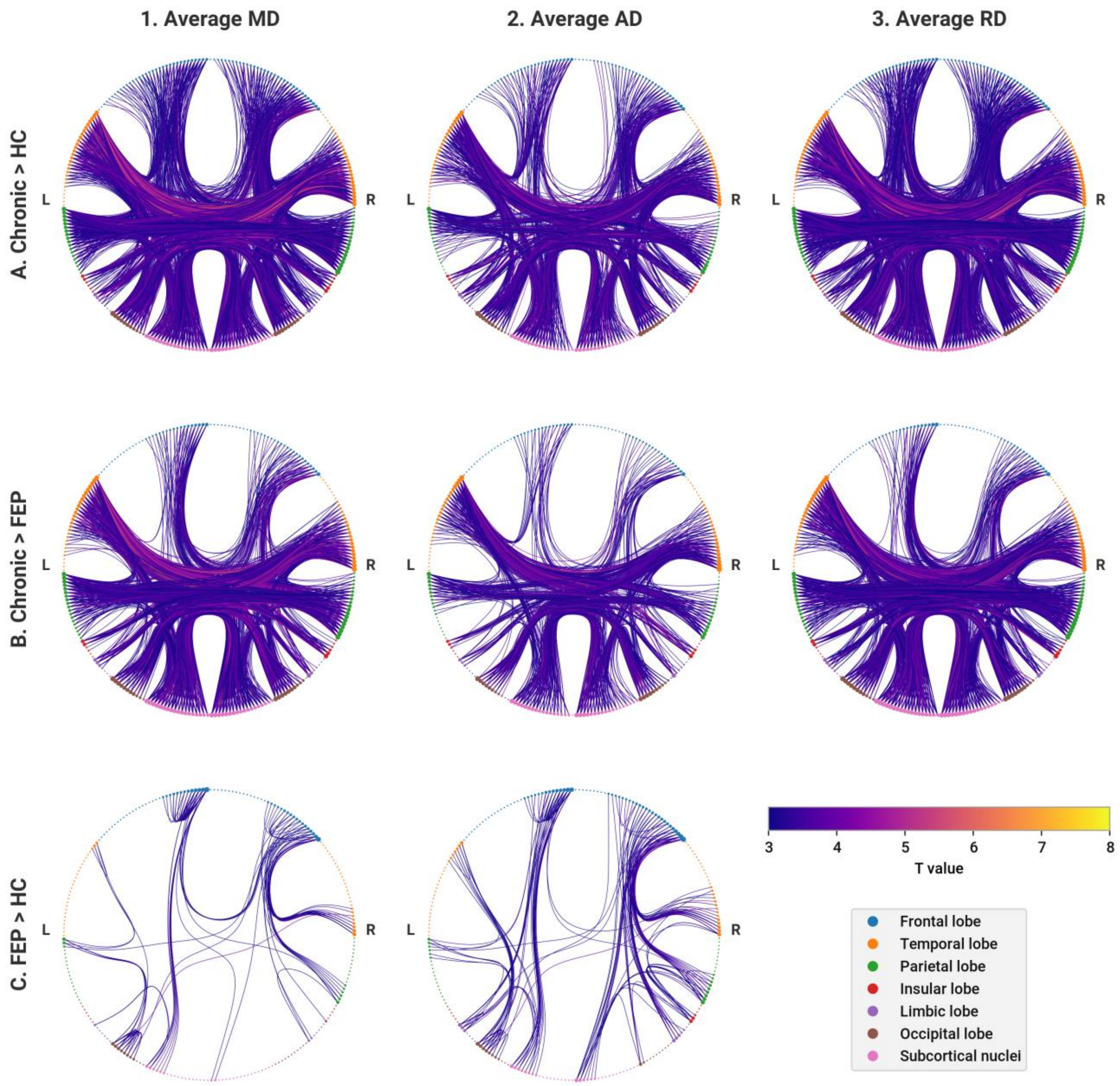


*Figure S2: AD, RD, and MD are significantly increased in FEP and chronic patients. For each network diagram, lines correspond to edges with a significant increase of the corresponding metric in the corresponding group. Nodes are colored according to brain region and sized according to number of connecting edges. Left hemisphere nodes are on the left side of each diagram, right hemisphere on the right. Rows: A. Edges of chronic patients with increased parameter values compared to HC. B. Chronic compared to FEP. C. FEP compared to HC. Columns: 1. Increased average mean diffusivity. 2. Increased average axial diffusivity. 3. Increased average radial diffusivity.*

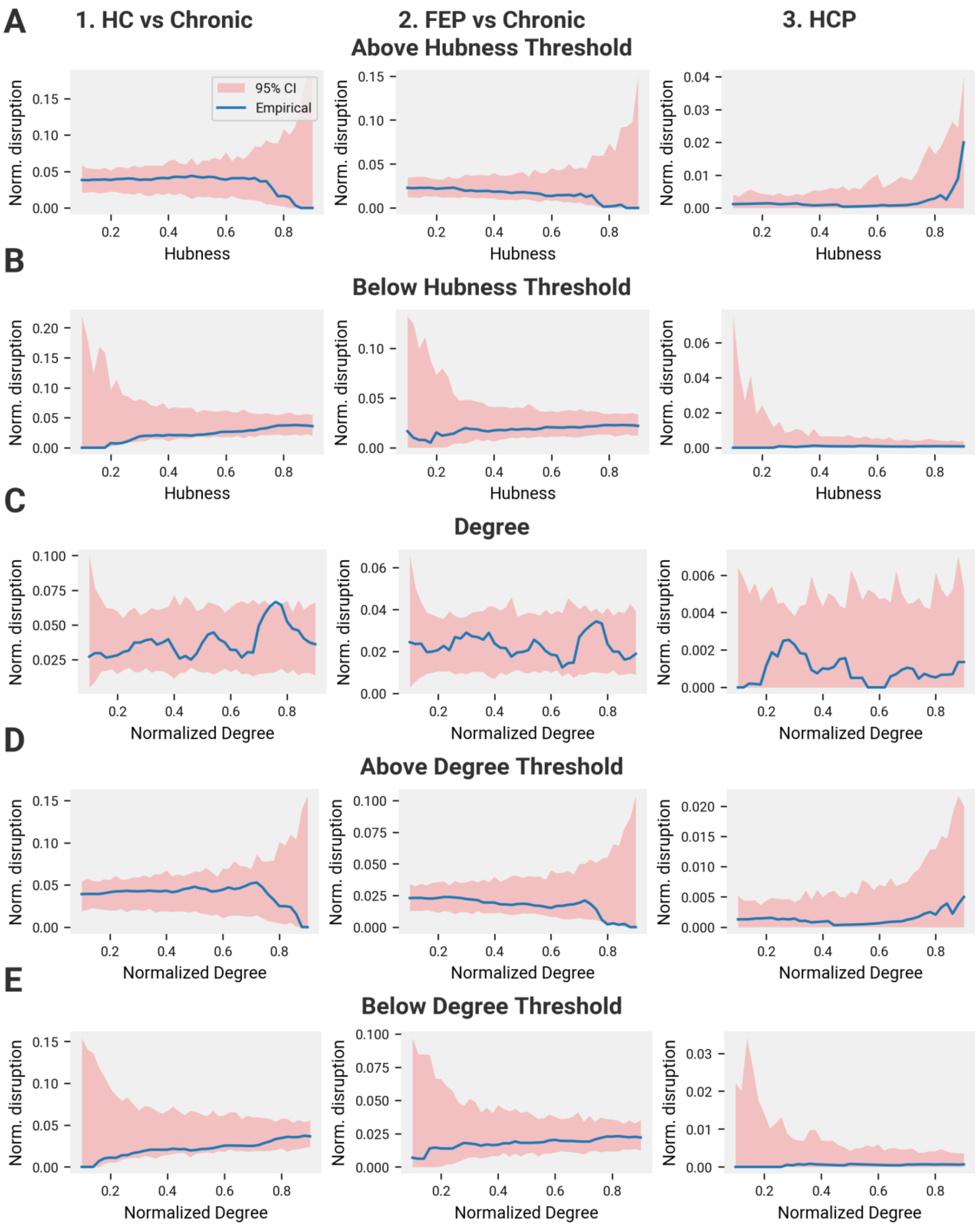


*Figure S3 (preceding page): Alternative measurements of disruption topology fail to find patterns. Charts represent the predominance of edges with significantly reduced FA in various subgraphs. In all rows, the left-most column represents data from the HC-chronic comparison, the center the FEP-chronic comparison, and the third the HC-EP comparison from the HCP-EP*

*dataset. The red shaded regions represents the 95% confidence interval of 10,000 random permutations (re-analyzed at each threshold value). For each permutation, a random subgraph equal to the size of the empirical subgraph of the respective threshold value is analyzed. A. X-axis charts a sliding lower-bound threshold of hubness. At each threshold, the proportion of edges disrupted in the subgraph of nodes with hubness higher than the threshold is graphed on the y axis. B. As in A, but the X-axis represents a sliding upper-bound threshold. C. As in Figure 3, the X-axis represents a sliding window. For each hubness value* $x$*, the proportion of edges with significantly lower FA from the entire graph that connect to nodes with hubness equal to* $x \pm 0.05$ *is calculated. Degree rank is used as the hubness measurement instead of the composite hubness score. D. As in A, but with degree rank. E. As in B, but with degree rank.*

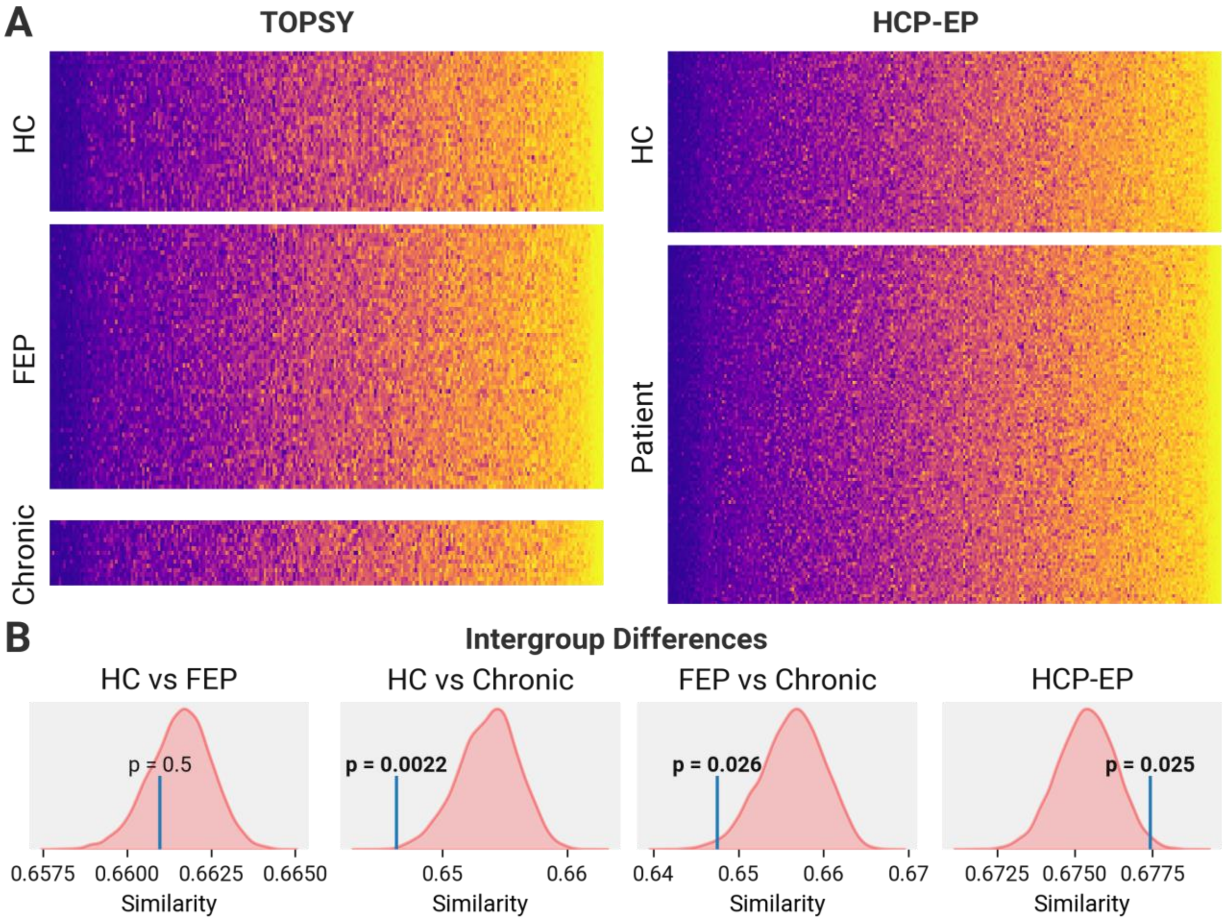


*Figure S4: Extended data showing comparisons of node hierarchy. A. Node hubness rankings for individual subjects. Nodes are rank-ordered along the x-axis according to increasing average hubness across all subjects. The left column shows TOPSY data, the right shows HCP-EP data. Each diagram is split into sections corresponding to the different patient subgroups. The relative ordering of nodes along the x-axis differs between the two datasets, but remains consistent within each dataset across all groups. B. Results of permutation analyses comparing the similarity of hubness rank-order lists between groups. Left most graph shows HC vs FEP, the next shows HC vs chronic, the next shows FEP vs chronic, and the rightmost compares the two HCP-EP groups. The red curve shows the random distribution obtained by 10,000 permutations, randomly shuffling participants between the two groups under consideration. The blue line shows the location of the emperical value. Significant p-values are displayed in bold.*

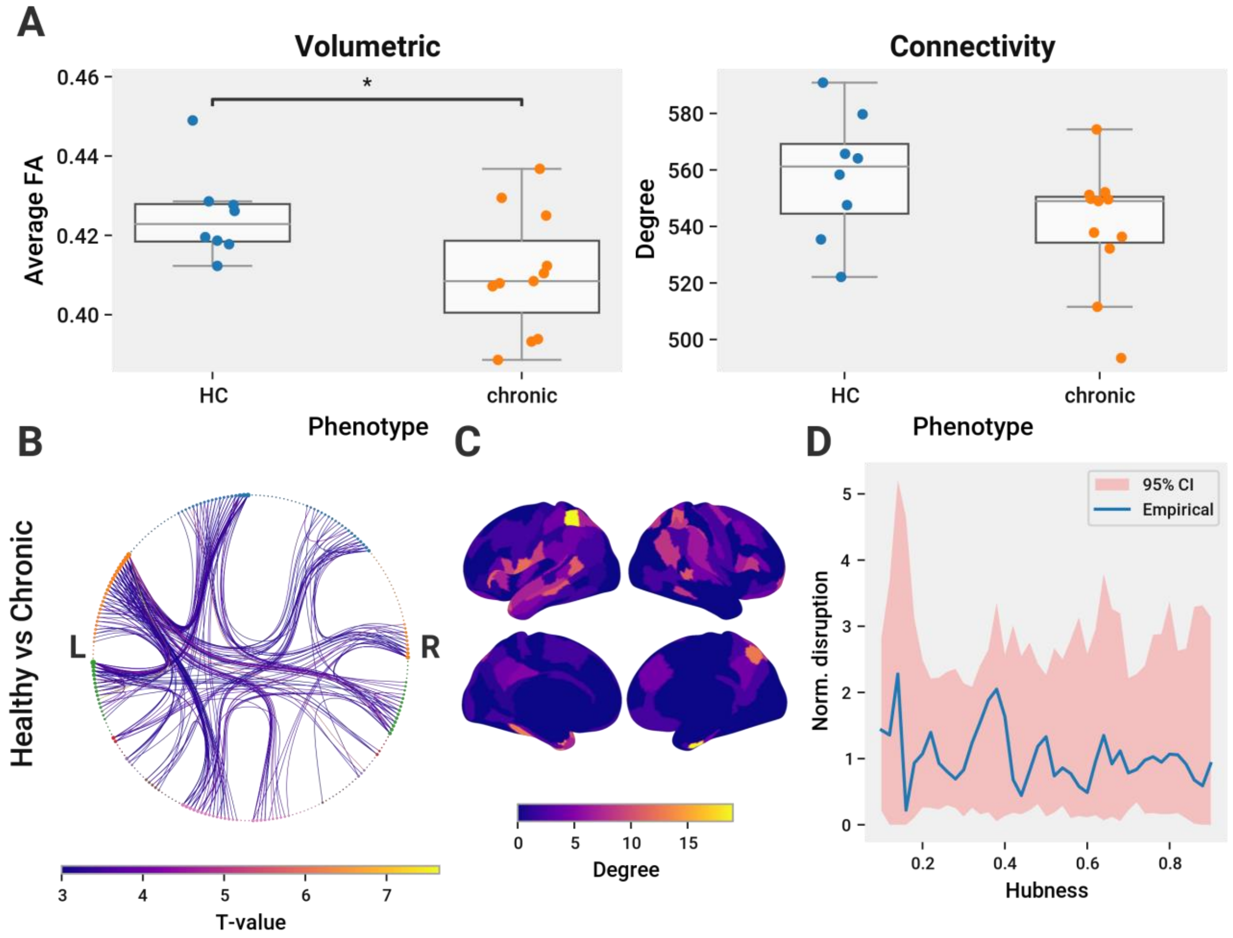


*Figure S5: Age-matched comparisons between HCs and chronic patients. A. Left: average FA across all connections is significantly lower in chronic patients than age-matched controls ($t(17) = 2.29, p = 0.035$). Right: average degree of the Brainnetome-parcellated connectome weighted with the logarithm of the SIFT2-weighted streamline count is not significantly lower in chronic patients. B. Edges with significantly lower FA in chronic patients compared to age-matched controls. Nodes from the Brainnetome parcellation are colored according to their brain region and sized according to the number of disrupted edges they connect to. The legend for brain regions is displayed in Figure 3. Edges are coloured according to their T-value. C. Cortical regions colored according to the number of disrupted edges connected. D. Topological organization of disrupted edges, as described in Figure 3 C.*

#### 2 SUPPLEMENTARY TABLES

*Table S1: Summary of significant voxels in Tract-Based Spatial Statistics (TBSS) experiments. Extent corresponds to the ratio of significant voxels to the total number of voxels in the skeletal mask, expressed as a percentage. Affected regions were determined by overlap between significant voxels and labelled regions in the ICBM-DTI-81 white matter atlas.[1] Superficial White Matter (SWM) affected regions were determined by visual inspection. Abbreviations: FA=Fractional Anisotropy; MD=Mean Diffusivity; RD=Radial Diffusivity; AD=Axial Diffusivity; Pos=Posterior; Ant=Anterior; Sup=Superior; Sup. Longt. Fasc.=Superior Longitudinal Fasciculus,*

| Model | Parameter | Extent (% Voxels) | Volume ($mm^3$) | Hemisphere | Affected Regions (ICBM-DTI-81) |
|---|---|---|---|---|---|
| HC > Chronic | FA | 0.863 | 1000 | Left | Body and Splenium of Corpus Callosum, Pos. Corona Radiata |
| Chronic > HC | MD | 8.82 | 10216 | Bilateral | Body, Splenium, and Genu of Corpus Callosum, Sup. and Pos. Corona Radiata, Pos. Thalamic Radiation, Sagittal Stratum, Sup Longt. Fasc. Tapetum |
| | | | | Left | Retrolentricular Limb of Internal Capsule, Ant. Corona Radiata, Cingulum |
| | | | | Right | Tapetum |
| | | | | SWM | Bilateral Parietal, Left Frontal, Right Occipital |
| Chronic > HC | RD | 11.1 | 12800 | Bilateral | Body and Splenium of Corpus Callosum, Retrolentricular Limb of Internal Capsule, Sup. and Pos. Corona Radiata, Pos. Thalamic Radiation, Sagittal Stratum, Sup. Longt. Fasc., Tapetum |
| | | | | Left | Cererbral Peduncle, Ant. And Pos. Limb of Internal Capsule, Cingulum |

| Model | Parameter | Extent (% Voxels) | Volume ($mm^3$) | Hemisphere | Affected Regions (ICBM-DTI-81) |
|---|---|---|---|---|---|
| | | | | SWM | Bilateral Parietal, Temporal, Occipital |
| FEP > Chronic | FA | | 1952 | Bilateral | Body and Splenium of Corpus Callosum, Sup. Longt. Fasc. |
| | | | | Left | Sup. and Pos. Corona Radiata, Cingulum, Tapetum |
| Chronic > FEP | MD | 0.083 | 96 | Right | Splenium, Post. Corona Radiata |
| Chronic > FEP | RD | 6.57 | 7608 | Bilateral | Body and Splenium of the Corpus Callosum, Sup. and Pos. Corona Radiata, Pos. Thalamic Radiation, Sup. Longt. Fasc., Tapetum |
| | | | | Left | Retrolentricular Limb of Internal Capsule, Cingulum |
| | | | | Right | Sagittal Statum |
| | | | | SWM | Bilateral Parietal, Right Temporal, Occipital |
| FEP > HC | MD | 5.41 | 6272 | Bilateral | Genu of Corpus Callosum, Ant. Corona Radiata, Sup. Longt. Fasciculus |
| | | | | Left | Pos. Thalamic Radiation |
| | | | | Right | Ant. Limb of Internal Capsule, Sup. Corona Radiata |
| | | | | SWM | Right Frontal, Left Occipital |
| FEP > HC | AD | 0.463 | 536 | Left | Genu of Corpus Callosum, Ant. Limb of Internal Capsule, Ant. Corona Radiata, External Capsule |
| | | | | SWM | Left Frontal |

*Table S2: Demographics of healthy control cohort age-matched to chronic patients.*

| | HC (n=8) | chronic (n=11) | HC vs chronic |
|---|---|---|---|
| Sex (M/F) | 6/2 | 9/2 | $\chi^2(1) = 0$, P = 1 |
| Age | 26.50 (2.07) | 31.00 (7.71) | t(17) = -1.6, P = 0.13 |
| Handedness (R/L) | 8/0 | 10/1 | $\chi^2(1) = 0$, P = 1 |
| Education | 15.50 (2.14) | 12.55 (2.42) | t(17) = 2.75, **P = 0.014** |
| SES | 3.75 (1.04) | 3.55 (1.29) | t(17) = 0.369, P = 0.72 |
| CAST | 6.00 (0.00) | 9.40 (5.82) | t(16) = -1.64, P = 0.12 |
| AUDIT-C | 3.00 (1.60) | 3.30 (2.58) | t(16) = -0.286, P = 0.78 |
| Smoker (yes/no) | 0/8 | 5/6 | $\chi^2(1) = 2.87$, P = 0.09 |
| Cannabis (yes/no) | 4/4 | 3/7 | $\chi^2(1) = 0.143$, P = 0.71 |
| SOFAS | 82.20 (2.59) | 57.09 (9.63) | t(14) = 5.64, **P = 6.1e-05** |
| PANSS-8 Total | 8.00 (0.00) | 15.18 (7.44) | t(17) = -2.71, **P = 0.015** |
| PANSS-8 Positive | 3.00 (0.00) | 7.18 (3.92) | t(17) = -2.99, **P = 0.0082** |
| PANSS-8 Negative | 3.00 (0.00) | 4.09 (1.76) | t(17) = -1.74, P = 0.1 |
| PANSS-8 General | 2.00 (0.00) | 4.00 (2.76) | t(17) = -2.04, P = 0.058 |
